# Intrinsic specificity differences between transcription factor paralogs partly explain their differential in vivo binding

**DOI:** 10.1101/208561

**Authors:** Ning Shen, Jingkang Zhao, Joshua Schipper, Yuning Zhang, Tristan Bepler, Dan Leehr, John Bradley, John Horton, Hilmar Lapp, Raluca Gordan

## Abstract

Members of transcription factor (TF) families, i.e. paralogous TFs, are oftentimes reported to have identical DNA-binding motifs, despite the fact that they perform distinct regulatory functions in the cell. Differential genomic targeting by paralogous TFs is generally assumed to be due to interactions with protein cofactors or the chromatin environment. Contrary to previous assumptions, we find that paralogous TFs have different intrinsic preferences for DNA, not captured by current motif models, and these differences partly explain differential genomic binding and functional specificity. Our finding was possible due to a unique combination of carefully designed high-throughput assays and rigorous computation modeling, integrated into a unified framework called iMADS. We used iMADS to quantify, model, and analyze specificity differences between 11 paralogous TFs from 4 distinct human TF families. Our finding of differential specificity between closely related TFs has important implications for the interpretation of the regulatory effects of non-coding genetic variants.

## INTRODUCTION

Transcription factors (TFs) interact with DNA in a sequence-specific manner, and these interactions represent a key mechanism in the regulation of gene expression. In eukaryotes, most TF-coding genes have undergone gene duplication and divergence during evolution^8–15^, resulting in many TFs having highly similar DNA binding domains and recognizing similar DNA sequence motifs. TFs with such properties that also belong to the same species are called ‘paralogous’ TFs. Some paralogous TFs have partly (or completely) redundant functions. Most mammalian TFs, however, have evolved regulatory functions that are distinct from their paralogs in the cell ^10, 11, 16^. For example, in the Class I subfamily of ETS transcription factors^18–20^, protein Ets1 is involved in B-cell, T-cell and natural killer cell differentiation^21–26^, while protein Elk1 regulates chromatin remodeling, serum response factor-dependent transcription, and neuronal differentiation^21,27–33^. In general, paralogous TFs accomplish a wide variety of independent or complementary molecular functions to regulate cellular phenotypes.

Many methods have been developed to learn TF-DNA binding specificity models from high-throughput *in vivo* and *in vitro* experimental data, ranging from simple position weight matrices (PWMs) to state-of-art deep learning models^34–40^. According to these models, paralogous TFs, especially the ones with high amino acid identity in their DNA-binding domains, tend to have indistinguishable DNA-binding specificities^34^. As an important consequence, this restricts the inference of TF-DNA interactions to family-wide predictions, rather than predictions for individual family members.

Since paralogous TFs are often co-expressed in the same cells but they perform different, sometimes even opposite, biological functions, being able to identify genomic binding sites of individual TF family members is critical. For example, while c-Myc is a well-known oncoprotein that promotes transcriptional amplification, its co-expressed paralog Mad is a tumor suppressor and represses gene expression^41–43^. Currently, little is known about the mechanisms that explain the differential genomic targets of paralogous TFs. Furthermore, when analyzing *in vivo* TF-DNA binding data, such as data from chromatin immunoprecipitation coupled with sequencing (ChIP-seq) assays^44^, many genomic studies do not even take into account the presence of paralogous TFs, or the fact TF family members present in a cell are likely to influence each other’s binding to the genome. Overall, given that the most mammalian TFs are part of large protein families with multiple paralogs expressed at the same time, it is surprising how little we know about how paralogous TFs achieve their unique specificities in the cell.

Here, we show that despite having similar DNA-binding domains, paralogous TFs have different intrinsic DNA-binding preferences and this explains, in part, their differential *in vivo* binding and functional specificity. We focus on closely related TFs reported to have indistinguishable DNA-binding motifs, but distinct sets of targets *in vivo*. We design custom DNA libraries containing putative TF binding sites in their native genomic sequence context, and we use *in vitro* genomic-context protein-binding microarray (gcPBM) assays^45^ to quantitatively measure binding of each TF to the genomic sequences in our custom library. The quantitative, high-throughput gcPBM measurements revealed extensive differences in binding specificity between most pairs of paralogous TFs tested in our study. Most differences are concentrated in the medium and low affinity ranges, which explains why there were missed by previous DNA-binding data and models (see Discussion). We emphasize that the differences observed in our study are due to the intrinsic DNA recognition properties of the TFs, because our gcPBM assays measure binding of purified TF proteins to naked DNA sequences. Importantly, though, the differences we identified *in vitro* help explain, in part, the differential *in vivo* binding profiles of paralogous TFs.

To quantify the differences in specificity between TF paralogs, we developed an innovative approach that combines binding data for paralogous TFs with data from replicate experiments to derive weighted least square regression models of differential specificity. We integrate our high-throughput data and computational models into a general framework called iMADS (<u>i</u>ntegrative <u>M</u>odeling and <u>A</u>nalysis of <u>D</u>ifferential <u>S</u>pecificity), which we provide as a publicly available web tool (http://imads.genome.duke.edu). Using iMADS data and models, we show that genomic sites differentially preferred by TF paralogs have different sequence features and DNA shape profiles, and they are involved in distinct biological functions. Finally, applying iMADS models of differential specificity to the analysis of genetic variants provides novel insights into TF binding changes due to cancer somatic mutations.

## RESULTS

### 1. Closely related paralogous TFs bind differently to their genomic target sites *in vitro*

Paralogous TFs have similar DNA-binding domains (DBDs). However, their DBDs are not identical, and the amino acid sequences outside the DBD region are quite different. We hypothesized that these differences in protein sequence could lead to differences in DNA-binding specificity. To test this hypothesis, we focused on 11 closely related human TFs from 4 distinct structural families: basic helix-loop-helix (bHLH), E26 transformation-specific (ETS), E2 factor (E2F), and Runt-related transcription factors (RUNX). The factors were chosen based on: (i) availability of high-quality ChIP-seq data showing both overlapping and unique *in vivo* genomic targets for the paralogous TFs^40, 46^, and (ii) previous reports that the paralogous TFs have identical binding specificities^20, 34, 37, 47, 48^. Focusing on putative genomic target sites in their native DNA sequence context, we asked whether paralogous TFs have identical DNA-binding preferences, as expected from their indistinguishable position weight matrix (PWM) models trained on either *in vitro* (Fig. 1A) or *in vivo* (**Supplementary Fig. 1**) data. *In vitro* PWMs for human TFs are typically derived from high-resolution binding data for a large set of artificial or randomized DNA sequences (e.g. universal PBM^49^ or SELEX-seq^38^ data), while *in vivo* PWMs are derived from low-resolution data on binding to genomic DNA (e.g. ChIP-seq^44^ data). Our experimental approach, using gcPBM assays, is innovative in that we measure TF binding to genomic DNA sequences, i.e. sequences that the TFs also encounter in the cell, but at high-resolution and in a controlled environment. Thus, we take advantage of critical aspects of both *in vitro* and *in vivo* approaches.

**Figure 1.**
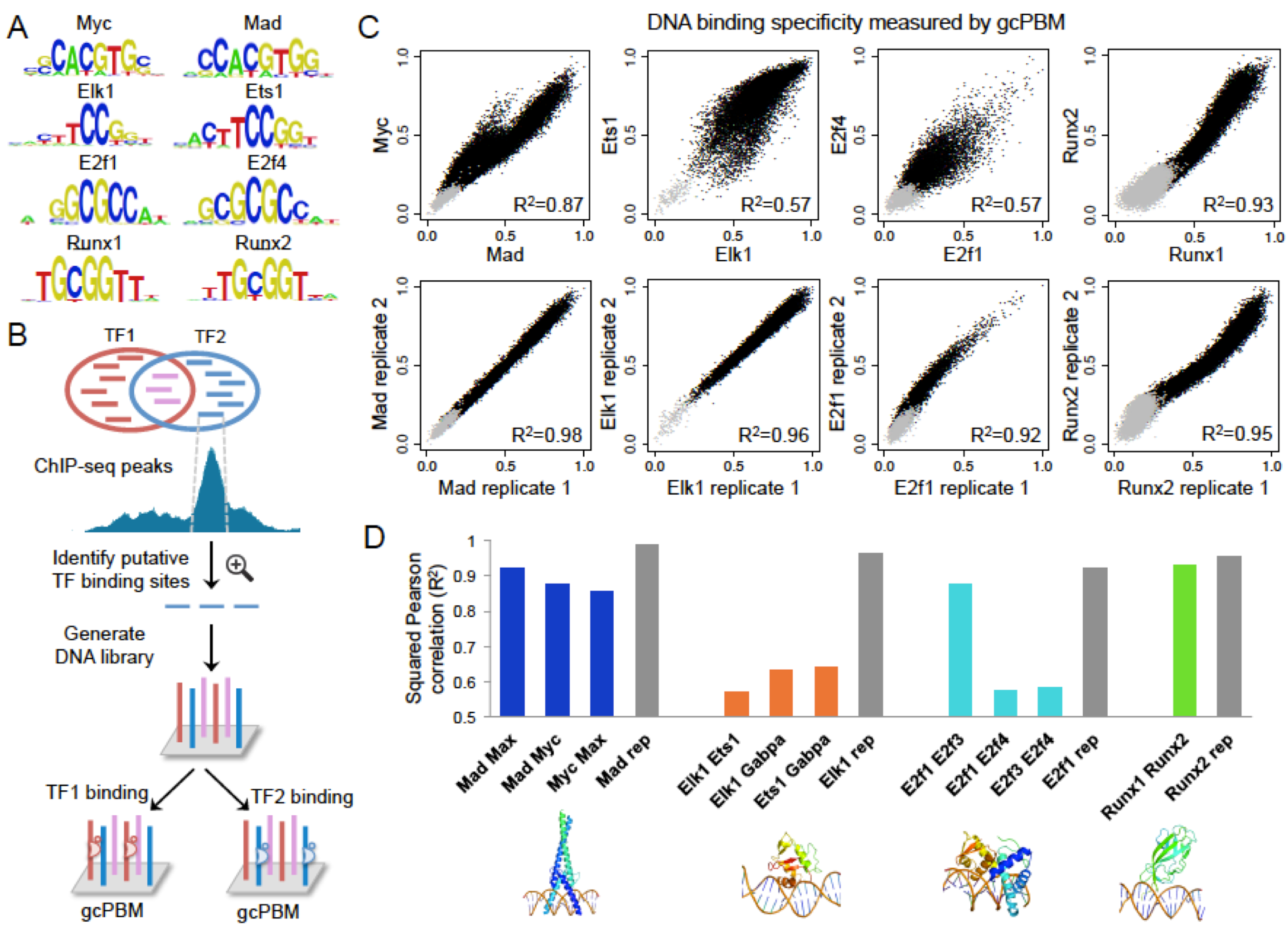
Paralogous TFs with indistinguishable PWMs show distinct *in vitro* specificities. **(A)** Examples of paralogous TF pairs with indistinguishable PWMs: Myc vs. Mad (bHLH family), Elk1 vs. Ets1 (ETS family), E2f1 vs. E2f4 (E2F family), Runx1 vs. Runx2 (RUNX family). **(B)** Design of gcPBM library containing putative binding sites for paralogous TFs, selected from unique (red and blue) and overlapping (purple) *in vivo* genomic targets, as identified by ChIP-seq. We used gcPBM assay to quantitatively measure *in vitro* binding of TF1 and TF2 to all selected genomic sites. **(C)** Direct comparisons of the binding specificities of paralogous TFs for genomic sites (top panels), in contrast to comparisons between replicate gcPBM data sets (bottom panels). Each point in the scatter plot corresponds to a 36bp genomic region tested by gcPBM. Black points correspond to genomic sequences centered on putative TF binding sites. Grey points correspond to negative control regions, i.e. sequences without binding sites. **(D)** Squared Pearson correlation coefficient (R^2^) for all pairs of paralogous TFs in our study (colored bars) and for representative replicate experiments (grey bars). Lower panel show example 3D structures for one TF in each family, as reported in the Protein Data Bank (PDB)^4^. From left to right: Mad:Max from bHLH family (PDB ID: 1NLW), Ets1 from ETS family (PDB ID: 2STT),E2F4:DP2 from E2F family (PDB ID: 1CF7), Runx1 from RUNX family (PDB ID: 1HJC).

The gcPBM assay measures the level of binding of a TF to tens of thousands of genomic regions simultaneously. Briefly, double-stranded DNA molecules attached to a glass slide (microarray) are incubated with an epitope-tagged TF. To detect the amount of TF bound to each DNA spot, the microarray is labeled with a fluorophore-conjugated antibody specific to the epitope tag, and scanned using a standard microarray scanner. The gcPBM protocol is similar to the universal PBM protocol of Berger and Bulyk^50^. The critical difference between the widely-used universal PBMs^20, 34–36, 49^ and the gcPBM assays in our study is in the design of the DNA library synthesized on the array. Universal PBMs use artificial sequences that cover all possible 10-bp DNA sites. Thus, they provide a comprehensive view of TF binding specificity for short sequences, but miss important information about the influence of flanking regions, which can significantly affect genomic binding^45^. In addition, universal PBMs suffer from significant spatial and location bias, due to the position of a probe on the microarray and the position of a TF binding site on the probe, respectively^49, 51^. In comparison, gcPBM libraries contain ~30,000 genomic sequences, 36-bp long, centered on putative binding sites for particular TFs or TF families (**Supplementary Fig. 2A**). Each sequence is represented six times in DNA spots randomly distributed across the array, and we use median values over replicate spots as TF binding specificity measurements. Our gcPBM libraries are carefully designed to: 1) capture the influence of flanking regions on TF-DNA binding by centering probes on the putative TF binding sites;2) minimize spatial bias by using median values over replicates spots; and 3) eliminate positional bias in the data by fixing the position of TF binding sites within probes. These critical characteristics of the experimental design lead to TF binding measurements that are highly reproducible, cover a wide range of binding affinities, and are in great agreement with independent binding affinity data (**Supplementary Fig. 2B,C**). In addition, as shown in our proof-of-concept study^45^, gcPBM measurements are sensitive enough to capture differences in specificity among related TFs.

To directly compare the binding specificities of TFs within each family, we designed family-specific DNA libraries containing putative binding sites in their native genomic context, and we used gcPBM assays to test *in vitro* binding of each family member to the selected genomic sites (Fig. 1B). The 11 TFs tested in our study are: c-Myc (henceforth referred to as Myc), Max, and Mxd1 (or Mad1, henceforth referred to as Mad) from the bHLH family; Ets1, Elk1, and Gabpa from the ETS family; E2f1, E2f3, and E2f4 from the E2F family; and Runx1 and Runx2 from the RUNX family. For all 11 TFs, the gcPBM assays provided quantitative measurements of *in vitro* specificity for tens of thousands of genomic sites. The vast majority of these sites were bound with affinities higher than negative controls, indicating that the selected genomic targets are specifically bound by the TFs in our study.

We analyzed the gcPBM data and, surprisingly, for most pairs of paralogous TFs we found extensive differences in their *in vitro* binding specificity for genomic sites, more than expected due to experimental noise (Fig. 1C,D). The way in which paralogous TFs differ is different for each family (Fig. 1C, top panels). bHLH proteins Mad and Myc bind similarly to many of their putative genomic targets, but there is a subset of sites bound with higher affinity by Myc than by Mad. ETS proteins Elk1 and Ets1 bind similarly to their high affinity genomic sites, but they diverge in specificity for medium and low affinity sites. E2F proteins E2f1 an E2f4 show differences across the entire affinity range, but mostly in the medium affinity sites. We note that although single gcPBM assays do not directly provide affinity measurements, the DNA-binding intensities measured by PBM do correlate very well with independently measured affinities (**Supplementary Fig. 2C**). To our knowledge, our data are the first to show that these closely related TFs have different intrinsic specificities for their genomic target sites.

The pair of paralogous TFs most similar to each other is Runx1 vs. Runx2 (rightmost panels in Fig. 1C,D). These factors act in different tissue types and are not typically co-expressed under normal cellular conditions^52–57^. Thus, their differential *in vivo* binding may be regulated through differential chromatin accessibility or different protein cofactors expressed in each cell type. The high similarity between the binding specificities of Runx1 and Runx2 is also not surprising given that their DNA-binding domains (DBDs) are 91% identical. Interestingly, considering all 11 pairs of paralogous TFs in our study, we did not see a correlation between DBD amino acid identity and similarity in DNA binding specificity (R^2^=0.01, **Supplementary Fig. 3**), indicating that amino acid identity might not be predictive for whether paralogous TFs prefer similar DNA target sites.

Overall, our gcPBM data show that most TF pairs converge in their specificity for high affinity sites, but bind differently to low and medium sites. This explains why many previous studies reported indistinguishable PWMs for these paralogous TFs^34, 37^: PWMs are best at capturing high affinity sites^58^, which are indeed bound the same way by the paralogous factors. However, medium and low affinity TF binding sites, which can play important regulatory roles in the cell^58–62^, are oftentimes bound differently by TF family members, and may contribute to the differential genomic binding and functional specificity of closely-related TFs. Therefore, we emphasize that quantitative measurements of TF binding over a broad affinity range, such as measurements provided by gcPBM assays, are critical for comparisons of paralogous TFs.

### 2. Generalizing TF binding specificities beyond the gcPBM measurements

In a single gcPBM experiment we can test up to ~30,000 genomic sites. However, a library of this size is still not sufficient to cover all putative genomic targets of human TFs. To generalize our TF-DNA binding measurements beyond the genomic sites tested on gcPBMs, we used ε-support vector regression (ε-SVR)^63^ to train positional k-mer regression models for all 11 TFs in our study. We used binary features to encode the identities of mononucleotides (1-mers), dinucleotides (2-mers), and trinucleotides (3-mers) at each position in the TF binding sites and their flanking regions (**Supplementary Fig. 4**), similar to our previous work^45,64^–^66^. A novel feature of the approach used in this work is that we build a ‘core-stratified’ SVR model for each TF, i.e. a separate SVR model is trained and tested for each ‘core motif of the TF (Fig. 2A). Core motifs are defined based on gcPBM data and, if available, based on prior structural knowledge about the interactions between DNA and TFs from each family (Methods). Core motifs are short (4-6bp) and capture the region within TF binding sites that has little degeneracy, likely because of direct interactions with residues in the DNA-binding domain of the TF (Fig. 2B,C). For example, for the bHLH transcription factor Mad, the core-stratified SVR model is based on five E-box or E-box-like cores (Fig. 2D).

**Figure 2.**
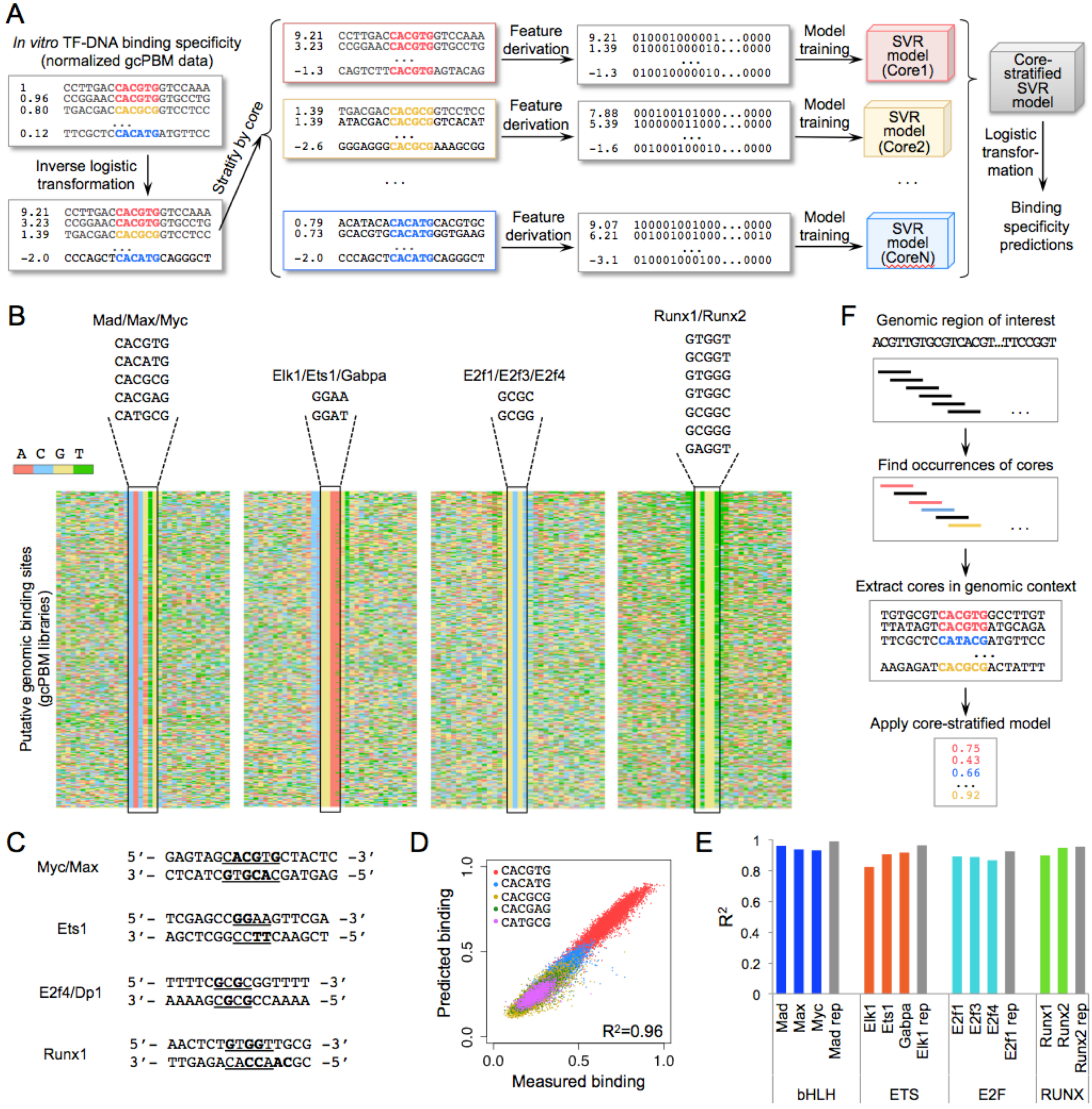
Core-stratified SVR approach to model TF-DNA binding specificity. **(A)** To build core-stratified SVR models of TF specificity, we start with normalized gcPBM data, apply an inverse logistic transformation, separate the gcPBM probes by their core motifs, derive features, and train one SVR model for each core. The predictions made by the SVR models are mapped back to a 0-1 range by applying the logistic transformation (Methods). **(B)** The core motif sequences for each group/family of TFs are shown. Heat maps show the DNA sequences for all gcPBM probes with binding intensity above the negative control range. **(C)** DNA bases that have direct interactions with TF residues, according to available X-ray crystal structures (PDB IDs: 1NKP for Myc/Max, 2STT for Ets1, 1CF7 for E2f4/Dp2, 1HJC for Runx1). DNA sequences tested in the crystal structure are shown. Core motifs are underlined, and bases that have direct interactions with protein residues are highlighted in bold. **(D)** Comparison of measured versus predicted DNA binding specificity for TF Mad (from embedded 5-fold cross-validation test; Methods). The cores used in the core-stratified SVR model are shown. **(E)** Prediction accuracy of core-stratified SVR models for all TFs used in this study, assessed as the squared Pearson correlation coefficient (R^2^) between measured and predicted DNA binding specificity (from embedded 5-fold cross-validation test; Methods). For comparison, gray bars show the correlation (R^2^) between replicate experiments. **(G)** Core-stratified SVR models can be used to make binding specificity predictions for any genomic region.

For all 11 TFs in our study, the core-stratified SVR models achieved high prediction accuracy (R^2^=0.82-0.96) on independent, held-out data, indicating that the models accurately capture TF-DNA binding specificity (Fig. 2E, **Supplementary Fig. 5**, **Supplementary Table 1**). All validations were performed using embedded 5-fold cross-validation tests (Methods). As baseline, we applied a nearest neighbor approach to the same folds as the core-stratified SVR. The nearest neighbor models had significantly lower accuracy, as expected (**Supplementary Fig. 6**). Given the high accuracy of our core-stratified SVR models, we can confidently use these models to predict TF binding to DNA sites not included in our gcPBM libraries. Importantly, we note that that core-stratified SVR models have prediction accuracy close to replicate experiments. Thus, while more complex models of specificity can be derived from our high quality gcPBM data, we do not expect such models to show large improvements in prediction accuracy compared to our core-stratified SVR models. Our simple, core-stratified approach is motivated by our observation that for different core motifs the sequences flanking the core contribute differently to the binding affinity (**Supplementary Fig. 7A,B**). To our knowledge, such interactions between the core TF binding motifs and the flanking base pairs have not been previously characterized in the literature. Training a separate SVR model for each core allows us to take into account the dependencies between the core motif and the flanking regions without resorting to complex computational models.

### 3. Modeling the differential DNA-binding specificity of paralogous TFs

Our gcPBM data revealed clear differences in the binding preferences of paralogous TFs for putative genomic target sites (Fig. 1). As with all high-throughput technologies that do not measure DNA-binding affinities directly, the binding measurements obtained by gcPBM are not directly comparable between TFs (one reason being that the samples used in experiments may have different concentrations of active TF protein). To address this shortcoming and perform a robust comparison between paralogous TFs, we developed and implemented a weighted regression approach (Fig. 3). As described below, this approach allows us to quantify specificity differences and to identify genomic sites differentially preferred by paralogous TFs.

**Figure 3.**
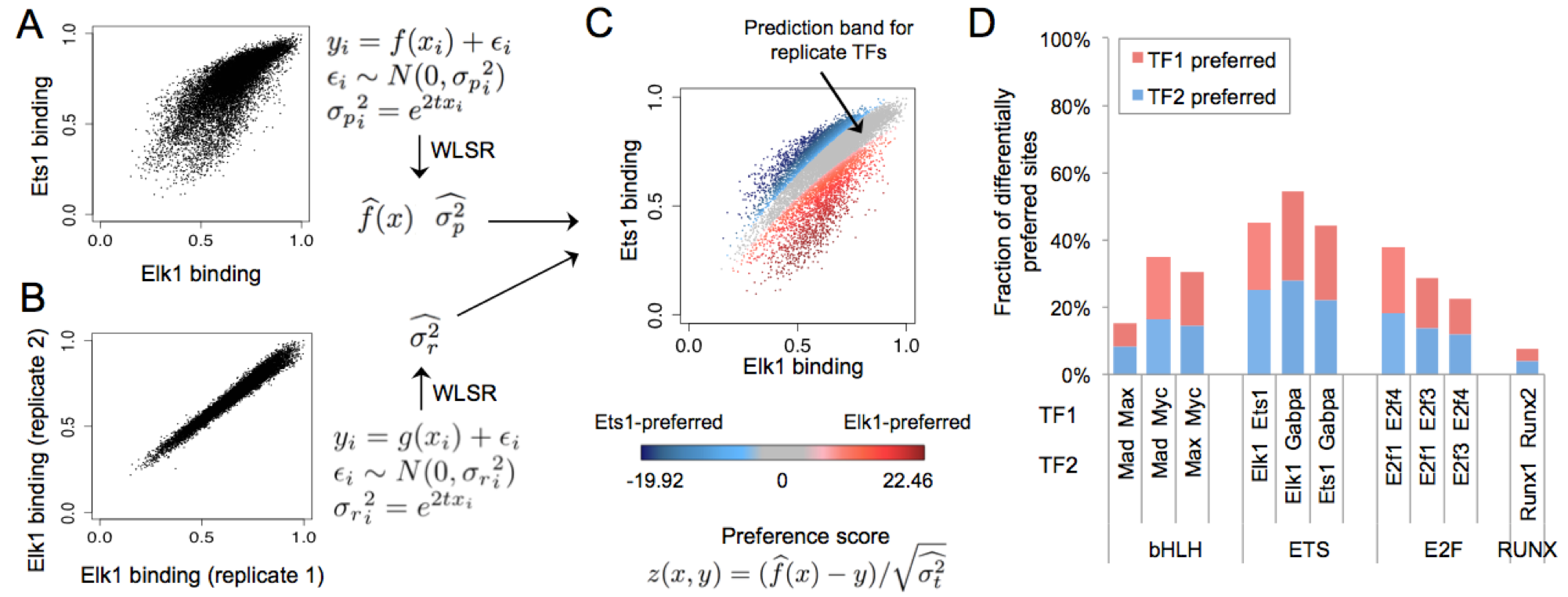
Modeling differential DNA-binding specificity. **(A)** Weighted least square regression (WLSR) is used to fit the gcPBM data of two paralogous TFs (here, Elk1 versus Ets1), and learn a linear or quadratic function 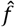, as well as the variance 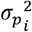 at every data point (i.e. genomic site) *i*. **(B)** Weighted least square regression (WLSR) is used to learn the variance structure for replicate gcPBM data sets. **(C)** By combining the variance learned from replicate data with the WLSR model for paralogous TFs, we compute a ‘99% prediction band for replicate TFs’ (grey), which contains genomic sites bound similarly by the two TFs (Methods). Genomic sites outside the prediction band are preferred by one of the two paralogous TFs (red: Elk1, blue: Ets1). The color intensity reflects the quantitative preference score computed according to the WLSR model. **(D)** Fraction of genomic sites, among the sites tested by gcPBM, which are differentially preferred by the paralogous TFs in our study.

Briefly, we apply weighted least square regression (WLSR) to fit the gcPBM data for two paralogous TFs (Fig. 3A), as well as replicate gcPBM data sets (Fig. 3B). Next, we integrate information about the variance learned from replicate data sets into the weighted regression model for the paralogous TFs, in order to calculate a ‘99% prediction band’ that comprises all genomic sites bound similarly by the two factors, i.e. sites for which the difference in binding specificity between TF1 and TF2 is within the noise expected for replicate experiments. Intuitively, one can interpret the 99% prediction band as follows: if TF1 and TF2 were replicates, then we would expect 99% of their target sites to fall within the prediction band. We consider the sites outside the prediction band as differentially preferred by TF1 versus TF2, and for each such site we compute a quantitative ‘preference score’ (Fig. 3C, Methods). We used our WLSR-based approach to compare all pairs of paralogous TFs in our study (**Supplementary Fig. 8**). For all TF pairs except Runx1 vs Runx2, we found that between 15% and 55% of the genomic sites tested by gcPBM were differentially preferred (Fig. 3D, **Supplementary Table 2**). Thus, our WLSR approach allows us to identify, for the first time, genomic sites differentially preferred by paralogous TFs, i.e. sites for which the difference in binding between TFs is larger than the variability observed in replicate experiments.

To facilitate the use of our WLSR models of differential specificity between paralogous TFs, as well as our core-stratified SVR models of binding specificity for individual TFs, we developed the iMADS web server: http://imads.genome.duke.edu. The web server allows users to apply our models for each TF or TF pair to make predictions on any genomic or custom DNA sequence.

### 4. Sequence and structural characteristics of genomic sites differentially preferred by paralogous TFs

We analyzed the differentially preferred genomic sites to determine sequence and structural features preferred by each TF. We found that the observed specificity differences between paralogous TFs are due both to the core binding site and the flanking regions, demonstrating the importance of including genomic flanks when measuring and comparing *in vitro* binding of these TFs. To identify significant differences in core and flanking preferences between TF1 and TF2, we applied the Mann-Whitney *U-* test to determine, for each core sequence and each 1-mer, 2-mer, and 3-mer feature in the flanking regions, whether the sequence feature is enriched in the set of TF1-preferred or TF2-preferred genomic sites. We report all p-values, adjusted using the Benjamini-Hochberg procedure, in **Supplementary Table 3.**

Core motifs play critical roles in TF-DNA recognition through direct interactions, mostly hydrogen bonds, between the proteins and DNA. This direct readout mechanism is a major contributor to the binding specificity of TFs^67^. In particular, direct readout in the core binding region is known to be different for different TF families^67^. Our results show that even *within* TF families, the core binding region can contribute to differences in binding specificity between factors (Fig. 4A, **Supplementary Fig. 9A,B**). For example, within the ETS family, the GGAT core is strongly preferred by Ets1 compared to both Elk1 (*p*=3.5×10^−99^, Fig. 4A) and Gabpa (*p*=1.8×10^−202^, **Supplementary Table 3**).

**Figure 4.**
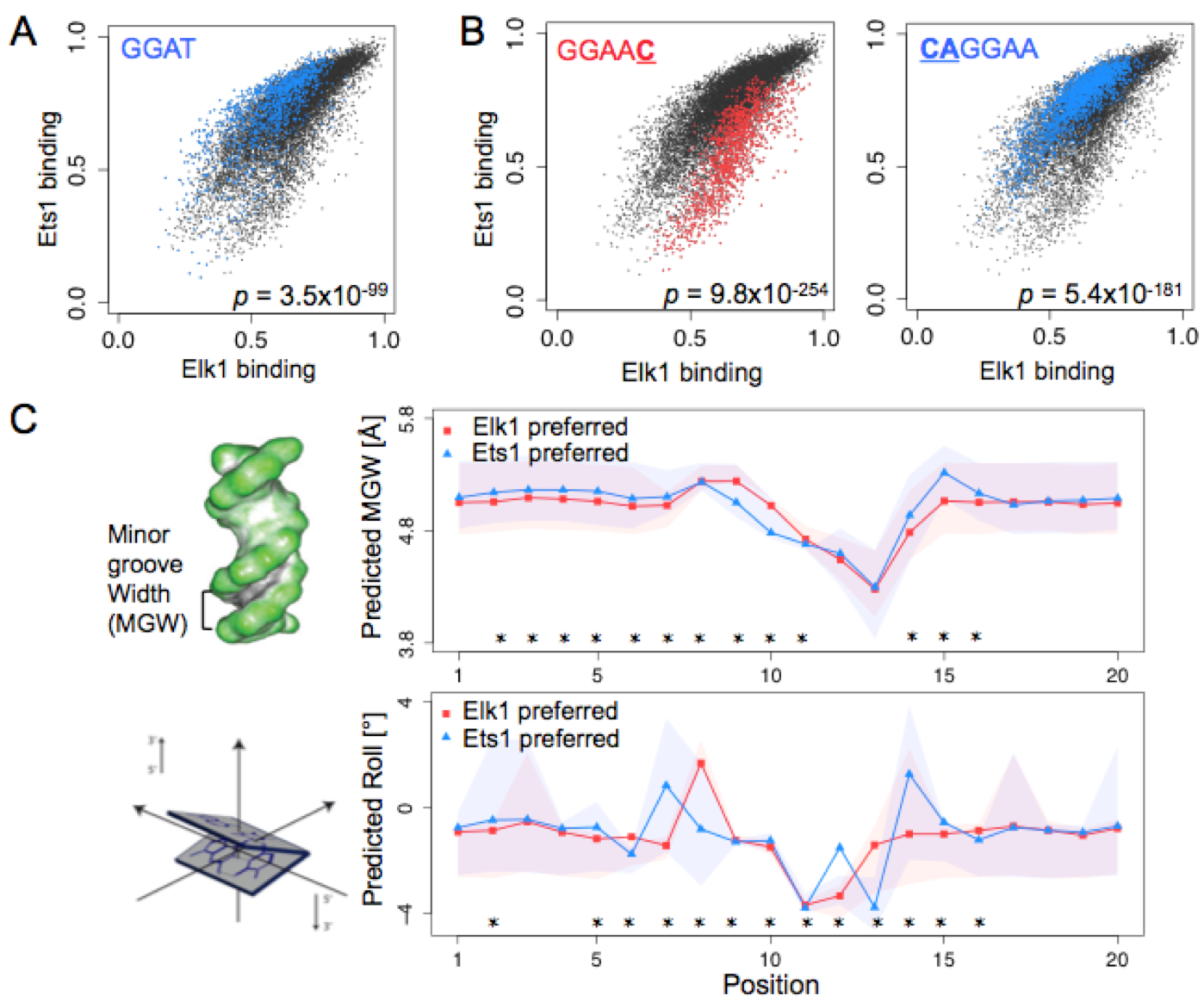
DNA sequence and shape preferences contribute to the differential specificity of paralogous TFs. **(A)**Core motif GGAT shows significant specificity preference for Ets1 vs. Elk1. *P*-value shows the enrichment of the core in Ets1-preferred sites, according to a Mann-Whitney *U*-test (Methods). **(B)** 1-mer and 2-mer sequences most differentially preferred by Elk1 or Ets1, among sites with the GGAA core. P-values were computed according to Mann-Whitney *U*-test (Methods). **(C)** Left: schematic of the minor groove width (MGW) and roll structural features. Right: MGW and roll profiles for genomic sites preferred by Elk1 versus Ets1. Stars (*) mark the positions within the binding sites (core or flanking region) that are significantly different between the two profiles (*p*-value < 10^−5^ according to Mann-Whitney *U*-test). Shaded regions show the 25^th^-75^th^ percentile ranges at each position.

Focusing on sequence features in the flanking regions, we identified numerous 1-mer, 2-mer, and 3-mer features differentially preferred by paralogous TFs (Fig. 4B, **Supplementary Fig. 9A,B, Supplementary Table 3**). For example, for the GGAA core bound by ETS factors, we found that C immediately downstream of the core is strongly preferred by Elk1, while CA immediately upstream of the core is strongly preferred by Ets1 (Fig. 4B). The differential preferences of paralogous TFs for flanking sequence features highlight the important role of genomic sequence context in establishing differential DNA binding between TF family members.

Flanking regions are likely contributing to TF-DNA binding specificity through indirect (i.e. shape) readout mechanisms. Using DNAshape^68^ predictions of minor groove width, roll, propeller twist, and helix twist, we found that paralogous TFs differ significantly in their preference for certain DNA shape features, especially for minor groove width and roll (Fig. 4C, **Supplementary Fig. 9C,D, Supplementary Table 4**). These findings are in agreement with previous hypotheses that DNA shape readout is often exploited to distinguish between TF family members^67^. Our data and models provide, for the first time, a way to comprehensively study the differences in DNA shape profiles preferred by paralogous TFs.

### 5. Differential *in vitro* specificity partly explains differential *in vivo* binding and functional specificity of paralogous TFs

To test whether the differences in intrinsic binding specificity between paralogous TFs, as observed in our gcPBM data, are relevant for differential *in vivo* binding, we applied iMADS models to make predictions of specificity and differential specificity on ChIP-seq peaks for Mad and Myc from H1SC cells, Elk1 and Ets1 from K562 cells, and E2f1 and E2f4 in K562 cells^46^. We selected these data sets because they were not included in the gcPBM design, and thus are completely independent of our training data. For each TF pair, we processed the ChIP-seq data to identify peaks for each TF, we merged the two lists of peaks, and for each peak we computed the natural logarithm of the ratio between TF1 and TF2 ChIP-seq signals (Methods). We then scanned the peak regions and used iMADS models to predict differential binding of TF1 vs. TF2 (e.g. Fig. 5A). To test whether the binding preferences captured by iMADS are relevant for the differential *in vivo* binding of paralogous TFs, we first performed a direct comparison between iMADS preference scores and differential ChIP-seq signal, computed for all peaks. As shown in Fig. 5B-D, the overall correlation is highly significant (*p*<10^−15^ for Mad vs. Myc and E2f1 vs. E2f4, p=0.003 for Elk1 vs. Ets1), although moderate (Spearman correlation *ρ*=0.28 for Mad vs. Myc, 0.44 for E2f1 vs. E2f4, and 0.06 for Elk1 vs. Ets1). Interestingly, this correlation is comparable or better than the correlation obtained for individual TFs by comparing their ChIP-seq data versus individual *in vitro* binding specificity models (*ρ*=0.14 for Mad, 0. 22 for Myc, 0.2 for E2f1, 0.26 for E2f4, 0.4 for Elk1, and 0.05 for Ets1; see Methods). These results were not surprising given that ChIP-seq data is noisy and contains numerous technical biases^69^, including formaldehyde crosslinking bias^70–75^, antibody specificity and variability problems^76–78^, technical artifacts due to highly expressed regions of the genome^79–81^, bias due to genome fragmentation and PCR amplification^82, 83^, etc. These biases can lead to false positive and false negative peaks, and they also significantly affect any quantitative estimates of *in vivo* TF binding levels derived from ChIP-seq data in ways that we do not understand well enough to correct^75^. It was also not surprising to find a lower correlation for the ETS proteins, compared to the bHLH and E2F proteins, given the low quality of the ChIP-seq data for Ets1 (Fig. 5E-G; **Supplementary Fig. 10**). Next, we binned the ChIP-seq data into 10 bins of equal size, after sorting the peaks in increasing order of the log ratio of TF1 vs. TF2 ChIP signal. We found significant differences between the distributions of iMADS preference scores in different bins, with higher iMADS scores corresponding, in general, to bins with higher log ratio of TF1 vs. TF2 ChIP signal (Fig. 5B-D, right panels). This was especially true for proteins with high-quality ChIP-seq data (Mad vs. Myc, and E2f1 vs. E2f4), as shown in Fig. 5B-C. Binning the ChIP-seq data averages out experimental noise and biases inherent to ChIP data sets^69–78, 82, 83^; at the same time, binning may average out important (although yet unknown) biological signals. The technical challenges of current ChIP-seq assays prevent a reliable quantification of how much of the iMADS variation in each bin is due to limitations of the *in vivo* data versus potential biological factors present in the cellular environment but missing in our *in vitro* system (see Discussion).

**Figure 5.**
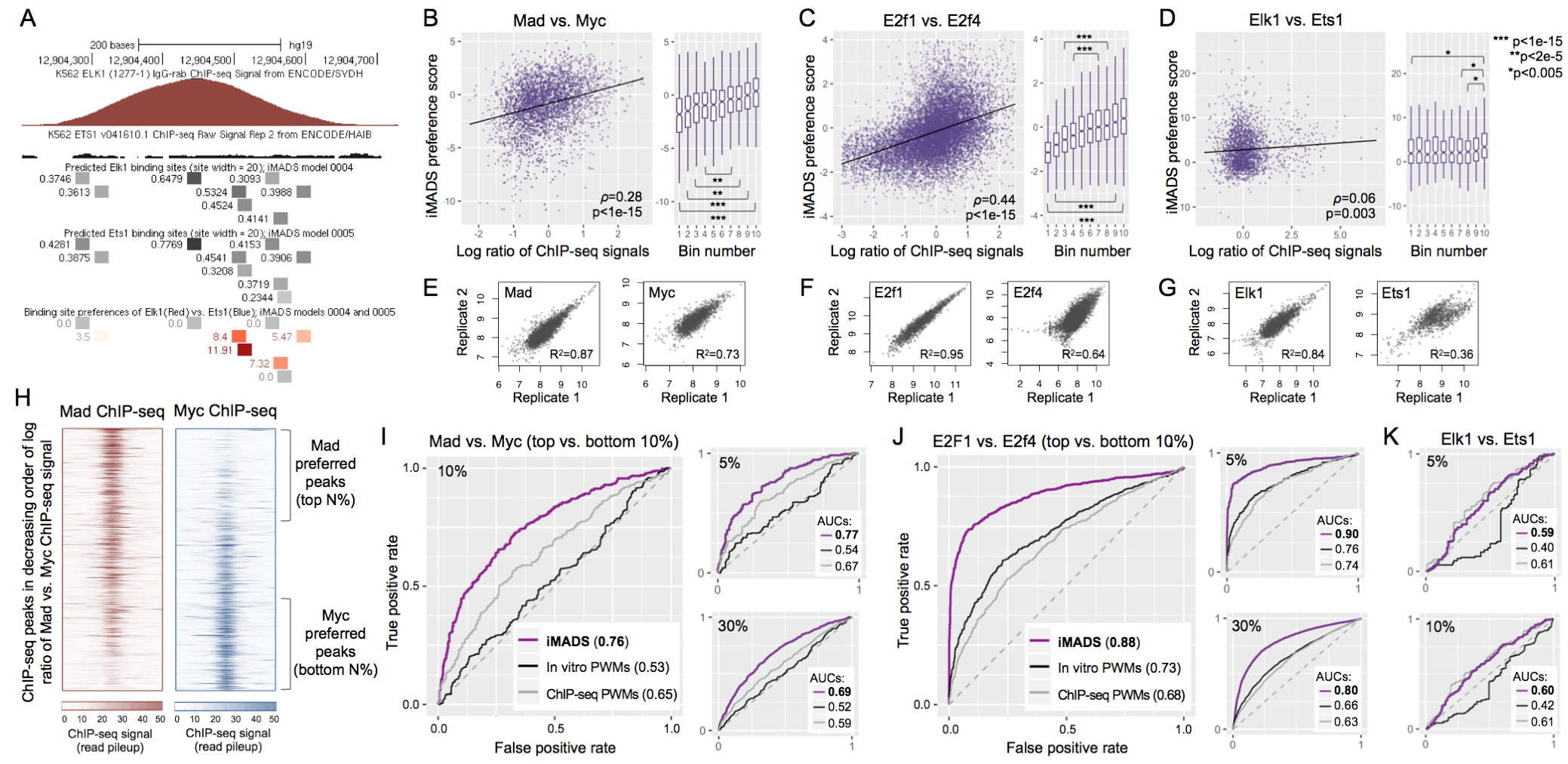
*In vitro* binding preferences of paralogous TFs partly explain their differential *in vivo* binding. **(A)** Genomic region bound *in vivo* by Elk1, but not Ets1 (according to ChIP-seq data^1^) contains binding sites with high preference for Elk1. **(B-D)** TF1 vs. TF2 *in vitro* binding preferences, as predicted using our iMADS preference models, have a significant correlation with *in vivo* binding preferences, as reflected by the log ratios of TF1 vs. TF2 ChIP-seq pileup signal (Methods). The Spearman correlation coefficient (ρ) and its statistical significance (p-value computed using the asymptotic *t* approximation^5^) is shown for each pair of TFs. Boxplots show the ChIP-seq data binned into 10 bins of equal size, based on the log ratios of TF1 vs. TF2 ChIP-seq signal. For each bin, the figure shows the median iMADS value and the 25th and 75th percentiles (lower and upper hinges); the whiskers extend from the hinges by 1.5 times the interquantile range. The distributions of iMADS preference scores in different bins were compared using Mann-Whitney *U-*tests. Due to outlier data points, the scatter plot for E2F factors is limited to peaks with log ratios in the [-3,2.5] interval, and iMADS preference scores in the [-4,4] interval. The full set of peaks, with log ratios in the [-7.79,5.41] interval and iMADS scores in the [-5.89,6.97] interval, are available in **Supplementary Table 5**. The full data sets (3,726 peaks for bHLH proteins, 13,004 peaks for E2F proteins, and 2,208 peaks for ETS proteins) were used to assess the correlations, to compute the best fit lines (shown in grey) and to compute the bins. **(E-G)** Pearson correlation coefficients between the ChIP-seq pileup signals computed from replicate ChIP-seq data sets. All data sets used in this analysis show good correlation, except for the Ets1 ChIP-seq data. Additional analyses of ChIP-seq data quality, performed using the ENCODE IDR^17^ pipeline, are shown in **Supplementary Fig. 10. (H)** ChIP-seq data for Mad and Myc, with peaks sorted in decreasing order of the log ratio of Mad vs. Myc signal. Regions of 1,000 bp centered at the peak summits are shown. The data can be used to identify ‘Mad-preferred’ and ‘Myc-preferred’ peaks, selected as the top and bottom N% of peaks, respectively. For different values of N, we tested how well iMADS models can distinguish between the peaks preferred by each TF. **(I)** Receiver operating characteristic (ROC) curves showing the performance of iMADS models of differential specificity, as well as PWM models trained on *in vitro* or *in vivo* data, in distinguishing Mad-preferred from Myc-preferred peaks. *In vitro* PWMs were derived from the same gcPBM data used to train iMADS preference models. *In vivo* PWMs were trained on the ChIP-seq data sets used for testing; thus, they are given an advantage over the other models. The area under the ROC curve (AUC) is shown for each model. AUC values vary between 0 and 1, with 0.5 corresponding to a random model. Results are shown for N=5, 10, and 30. **(J)** Similar to I, but for E2F proteins E2f1 versus E2f4. **(K)** Similar to I and J, but for ETS factors Elk1 and Ets1, and showing the results for N=5 and 10. Additional results for all three TF pairs are available in **Supplementary Table 6**.

To further analyze the relationship between TF binding *in vitro* and *in vivo,* we performed receiver operating characteristic (ROC) curve analyses, an approach that is widely used to evaluate the predictive power of DNA binding models with respect to *in vivo* ChIP-seq data^36, 39, 84–94^. Briefly, we aimed to evaluate how well our iMADS models of differential specificity can distinguish between TF1-preferred and TF2-preferred ChIP-seq peaks, defined as the top N% and bottom N% of peaks, respectively, sorted according to log ratio of ChIP signals (Fig. 5H). We found that our iMADS models perform remarkably well. For bHLH proteins Mad vs. Myc, the iMADS model achieved areas under the ROC curve (AUCs) of 0.69-0.77, depending on the fraction of peaks chosen for the classification test (top and bottom 5%-30% of peaks, Fig. 5I). In addition, the performance of iMADS models was superior to that of PWM models trained on either *in vitro* or *in vivo* data. In our comparisons we used *in vitro* PWMs trained on the same gcPBM data as the iMADS models. For the *in vivo*-derived PWMs, we used models trained on the same ChIP-seq data sets used for testing, thus giving *in vivo* PWMs an important advantage over the other models (see Methods). Nevertheless, our iMADS models of differential specificity performed best (Fig 5I), which is impressive given that they were trained on independent data and using independent genomic sequences. The performance of iMADS models was even better for E2F proteins E2f1 vs. E2f4, with AUCs between 0.8 and 0.9 (Fig. 5J). Even in the case of ETS proteins, which have poorer quality ChIP-seq data, the performance of our iMADS models (trained on independent data) was comparable to the performance of *in vivo* motifs (trained on the ChIP-seq data itself), especially when focusing only on the top vs. bottom 5-10% of the peaks (Fig. 5K). Thus, our results show the *in* vitro-derived iMADS models of differential specificity have significant predictive power *in vivo*.

Not surprisingly, we found that the correlation between iMADS preferences scores and differential ChIP-seq data is lower for TFs with low-quality ChIP-seq data, such as the ETS protein Ets1 (Fig. 5G). In such cases, since the ratios of ChIP-seq signal contain very little information, an alternative way to analyze *in vivo* binding is to focus only on the high-confidence peak regions that are unique to each TF. Such an analysis is presented in **Supplementary Fig. 11**, where we show that Elk1-unique peaks (i.e. genomic targets bound only by Elk1) contain sites with larger Elk1 preference scores, while Ets1-unique peaks contain sites with larger Ets1 preference scores. Thus, these results show that the differential specificity observed *in vitro* by gcPBM, and modeled using our iMADS framework, partly explains the differential *in vivo* genomic binding of paralogous TFs.

Paralogous TFs may achieve functional specificity through their differential DNA-binding specificity. To test this hypothesis, we focused on the genomic sequences in gcPBM data that are differentially preferred by paralogous TFs, and we compared the biological functions of genes in the neighborhood of these genomic binding sites. As shown in **Supplementary Fig. 12** for the example of Elk1 and Ets1, we successfully recovered different gene ontology (GO) terms that are enriched for genes associated with differentially preferred sites, with many of the terms reported previously in independent studies for the individual ETS factors^30, 95–101^. This suggests that sequences preferred differently by paralogous TFs *in vitro* are not only important for their differential genomic binding *in vivo*, but also help individual TFs achieve functional specificity in the cell.

To facilitate analysis of differential *in vivo* binding and functional specificity of paralogous TFs, users can easily access genome-wide predictions of TF binding specificity (from core-stratified SVR models; **Supplementary Fig. 13A,B**) and differential specificity (from WLSR models; **Supplementary Fig. 13C**) through the iMADS web server. In addition, users can focus on specific regions around genes, specify custom lists of genes or genomic coordinates to analyze, view predictions in the web server or in the UCSC genome browser, and make predictions of TF binding specificity and differential specificity for any DNA sequence of interest.

### 6. Disease-related genetic variants have differential effects on the specificity of paralogous TFs

Current studies of the effects of non-coding variants on TF-DNA binding focus on predicting changes in the DNA-binding specificity of individual TFs, assessed using simple PWMs^102–105^ or complex models^7, 39, 106^, but ignoring the fact that multiple paralogous factors are co-expressed in the cell and can influence each other’s binding to the genome. The iMADS models allow us to test, for the first time, whether non-coding variants/mutations have differential effects on the binding specificity of paralogous TFs.

To illustrate how one can use our iMADS models and web server to analyze non-coding variants, we use as a case study the somatic mutation rs786205688, associated with malignant prostate cancer^2^. The mutation resides in the POLK gene region, which is important for DNA damage repair, and it creates a binding site for the ETS family of TFs. According to current models, the newly created binding site has similar specificities for Ets1 and Elk1. However, according to the iMADS model of Elk1 versus Ets1 binding preference, the new site is highly preferred by Elk1, and bound only nonspecifically by Ets1, indicating that the functional effect of this mutation could be due to increased Elk1 binding (Fig. 6A). This hypothesis is consistent with the fact that up-regulation and activation of Elk1 has been reported to associate with malignancy of prostate cancer, and inhibition of Elk1 has been proven effective on inhibiting growth of prostate cancer cells^33^. In **Supplementary Fig. 14** we present the simple steps that users can follow to analyze non-coding variants, such as rs786205688, for their effect on binding of paralogous TFs. We note that the goal of such an analysis is not to conclusively identify a causal relationship between the variant and the phenotype, but to generate mechanistic hypotheses for follow-up analyses.

**Figure 6.**
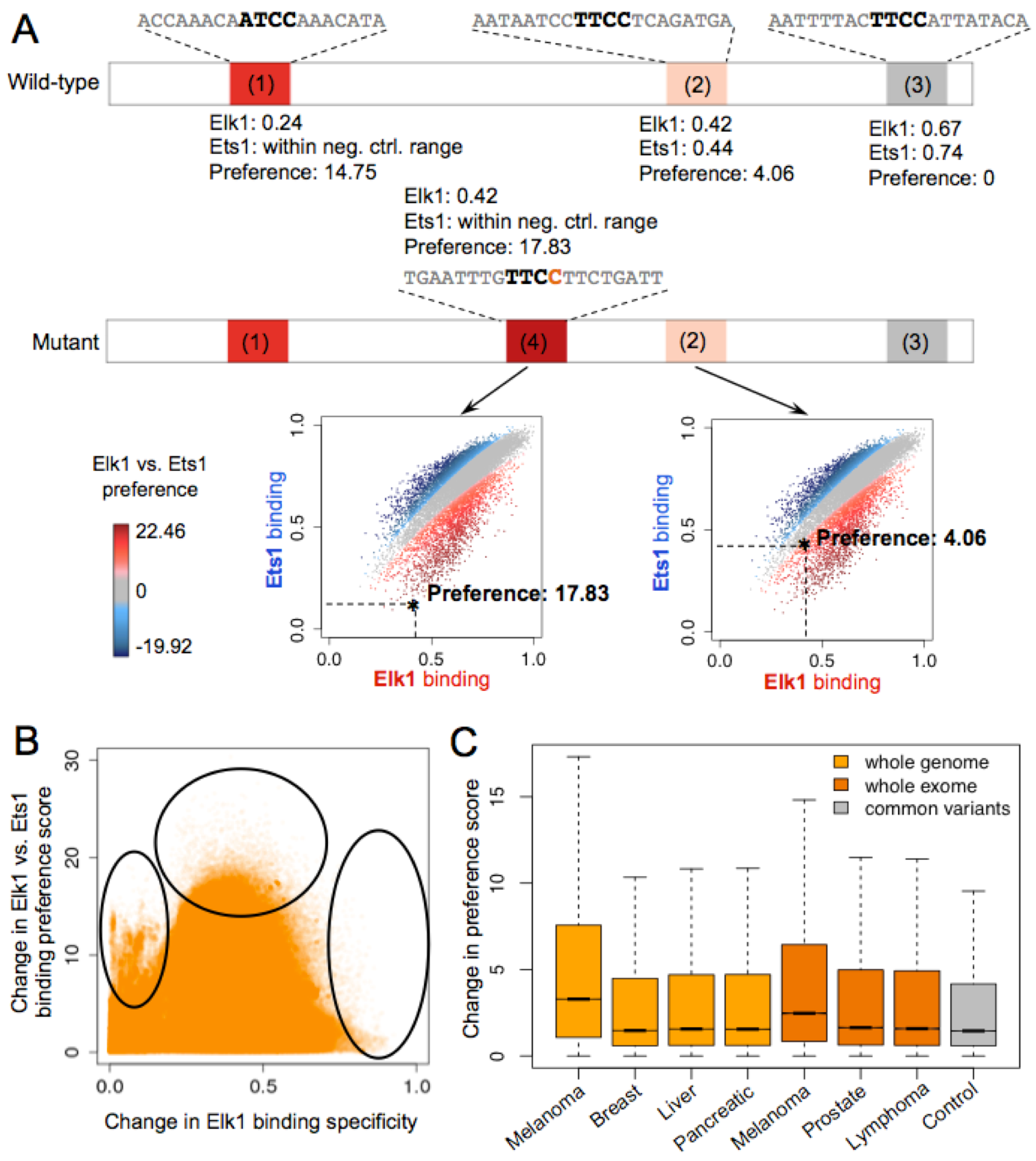
Analyses of non-coding somatic mutations using iMADS models of specificity and differential specificity. **(A)**Example of iMADS predictions for variant rs786205688, a somatic mutation in the POLK gene region associated with malignant prostate cancer^2^. Plots show predicted ETS sites in a 300-bp genomic region centered on the somatic mutation (chr5:74893909). Binding sites (1), (2), and (3) are present both in the wild-type (reference) and the mutant (tumor) sequences. Binding site (4) is created in the tumor by the somatic mutation. The color of the binding sites reflects the Elk1 vs. Ets1 preferences, as computed by iMADS. Scatterplots show the binding specificities and preferences of sites (4) and (2) (marked by stars) compared to the genomic sites tested by gcPBM in our study. **(B)** Scatterplot of the absolute change in Elk1 binding score compared to the absolute change in preference score of Elk1 vs. Ets1, for non-coding somatic mutations identified in melanoma cancer patients (ICGC data set SKCA-BR^3^). Ovals highlight sets of mutations discussed in the main text. **(C)** Boxplot comparing the changes in preference score between non-coding somatic mutations identified in different types of tumors^3^, versus a control set of non-coding variants from the 1000 Genomes Project^6,7^ The somatic variants were identified from either whole-genome (light orange bars) or whole-exome (dark orange bars) sequencing data from ICGC (Methods). The control variants (grey bar) were randomly selected among common variants with minor allele frequency > 0.01, as reported in the 1000 Genomes Project^6,7^ For all tumor types, preferences changes are significantly larger for somatic mutations than for common variants: one-sided Mann-Whitney U test p-value < 2.2e-16.

We also analyzed iMADS predictions for somatic mutations from melanoma whole-genome sequencing data^3^, to identify general patterns of TF preference change in disease-related mutations. Interestingly, we found a subset of mutations (Fig. 6B, left oval) that have almost no change in Elk1 binding specificity, but large changes in Elk1 vs. Ets1 preference. Such mutations would be completely ignored by analyses focusing only on changes in Elk1 binding specificity. In addition, mutations with the largest change in Elk1 specificity tend to have small changes in preference score (Fig. 6B, right oval). This was expected, as large changes in binding specificity will correspond to high affinity sites (in either wild-type or mutant sequences), which are bound similarly by the two TFs. Interestingly, we also found that mutations that maximize the change in Elk1 vs. Ets1 preference (Fig. 6B, top oval) are not the ones that maximize the change in Elk1 binding specificity. Thus, if one uses the change in Elk1 binding score to prioritize mutations, these mutations would not get high priority, although they could have large changes in Elk1 binding *in vivo* due to the change in preference relative to Ets1.

Next, we extended our analysis of non-coding somatic mutations to several tumor types with publicly available whole-genome or whole-exome sequencing data from the International Cancer Genome Consortium^3^. (We note that non-coding mutations in close proximity to coding regions can be identified from exome sequencing data.) We analyzed non-coding mutations from melanoma, breast cancer, liver cancer, pancreatic cancer, prostate cancer, and lymphoma, as well as a control set of common non-coding variants (Methods). We found that cancer mutations lead to significantly larger changes in Elk1-Ets1 preferences scores (Fig. 6C), suggesting that changes in the relative preferences of paralogous TFs could have important phenotypic effects. Interestingly, the changes in Elk1 vs. Ets1 binding preference were much more pronounced in melanoma compared to other tumor types. A major mutagenic factor in melanoma is ultraviolet light, which causes pyrimidine dimers, a type of DNA damage that occurs between neighboring pyrimidine nucleotides. As a result, melanoma samples have a very high frequency of C>T mutations, particularly at CC and TC dinucleotides. Given the high specificity of ETS family proteins for TTCC/GGAA and ATCC/GGAT cores, it is perhaps not surprising that binding of this family of TFs is more likely to be affected in melanoma than other tumor types. Our iMADS models of differential specificity allow us, for the first time, to study the effects of non-coding somatic mutations on the genomic binding of individual TF family members, taking into account the fact that a particular TF can be affected either directly (by mutations that change its specificity) or indirectly (by mutations that change the specificity of competing TF family members). To our knowledge, iMADS is the only available tool that allows users to analyze changes in relative preferences of paralogous TFs due to genetic variation, and we expect it to provide important insights into the interpretation of disease-causing non-coding mutations.

## DISCUSSION

DNA binding specificity is a fundamental characteristic of TFs. Nevertheless, the contribution of intrinsic sequence specificity to the differential *in vivo* binding of paralogous TFs is a largely unexplored area of research. Focusing on 11 paralogous TFs across 4 distinct protein families, we show that differences in intrinsic specificity, not captured by current DNA motifs model, can be critical for TF family members to distinguish between their genomic targets and achieve functional specificity in the cell. The integrated computational-experimental approach described in our study is general and can be applied to any pair of paralogous TFs.

Integrating quantitative measurements and modeling, as in our iMADS framework (**Supplementary Fig. 15**), is key to identifying and characterizing the differences between paralogous TFs. Most previous studies of TF-DNA binding focused on high affinity sites, which we find to be bound with similar specificities by paralogous TFs. For example, HT-SELEX/SELEX-seq assays37, 38, 107 exponentially enrich for high affinity sites, which limits their ability to obtain quantitative measurements for medium and low affinity sites. Our gcPBM data clearly shows that medium and low affinity sites have important contributions to the differential binding of paralogous TFs, in agreement with previous studies recognizing the importance of such sites for TF-DNA binding *in vitro* and *in vivo*^20, 35^. In addition, a critical feature of our experimental approach is that we tested TF binding to DNA sites in their native genomic sequence context, but in a controlled environment where we can directly assess and compare the intrinsic specificities of TFs for DNA. As shown by our results (Fig. 3, **Supplementary Fig. 9**), genomic sites differentially preferred by paralogous TFs have different sequence and shape features in their flanking regions. Importantly, we note that these contributions from the flanking regions are not due to cofactors or other influences, but to the intrinsic recognition properties of the paralogous factors.

Our results show that differences in intrinsic binding specificity between paralogous TFs have a significant contribution to differential *in vivo* binding (Fig. 5). Currently, this contribution cannot be reliably quantified using existing *in vivo* ChIP-seq data, as it is well known that traditional ChIP-seq methodologies are not inherently quantitative^108^. The frequently poor affinity, specificity, and reproducibility of antibodies represent a major source of experimental error in ChIP-seq^109^ (**Supplementary Fig. 16**). In addition, technical biases due to formaldehyde crosslinking^70–75^, highly expressed regions of the genome ^79–81^, genome fragmentation and PCR amplification^82, 83^ confound the true *in vivo* binding levels of paralogous TFs. When considering comparisons between ChIP-seq data sets, differences in experimental handling of the samples, as well as differential amplification prior to sequencer loading can also make direct ChIP-based comparisons problematic^109^. On the other hand, gcPBM assays provide direct measurements of TF binding, they have a simple design and few steps where technical biases could be introduced. But gcPBMs only capture DNA binding specificity coming directly from the protein, as the assays are performed using naked DNA and purified proteins. intermediate’ assays, such as sequencing-based assays that test TF binding to purified genomic DNA might be helpful, as they would have many of the same biases as ChIP-seq, but would lack any additional *in vivo* factors. Several modifications to traditional ChIP-seq protocols have also been proposed recently in order to allow quantitative comparisons between data sets^108–110^, primarily for histone proteins, by using spike-in controls. In addition, combining assays such as CUTandRUN^111^ or ChEC-seq^112^ (which alleviate some of the technical biases of ChIP-seq) with ChIP-exo^113^ or ChIP-nexus^114^ (which improve the resolution of the *in vivo* data), may improve our ability to quantify the *in vivo* binding levels of human paralogous TFs in the future. Currently though, for human TF proteins, traditional ChIP-seq is still the technique used by individual labs and research consortia, including the ENCODE consortium that generated the data used in our study^46^. Nevertheless, despite the shortcomings of current *in vivo* data, our results provide compelling evidence that differential intrinsic specificity of paralogous TFs, reported and quantified for the first time in our study, is likely one of the mechanisms for achieving differential genomic targeting in the cell.

Our discovery of the differential binding specificity between closely related TFs has major implications for interpreting the effects of non-coding genetic variants and mutations. Some disease-causing mutations could significantly affect binding of a TF by changing its preference relative to other family members expressed in the same cells. To our knowledge, no previous studies of non-coding genetic variations takes into account the potential influence of competing TF family members. Our analysis of somatic non-coding mutations shows that mutations that maximize the change in preference between paralogous TFs are not those that maximize change in specificity for either TF. This suggests that focusing on changes in binding specificity for individual TFs, as in previous studies, has limited power in understanding the effects of non-coding mutations on TF binding.

Given that the most mammalian TFs are part of large protein families with multiple TF paralogs expressed at the same time, it is surprising how little we know about how paralogous TFs achieve their unique specificities in the cell. We acknowledge that *in vivo* binding of a TF is a result of many complicated factors, including not only the intrinsic DNA-binding specificity of that TF and its paralogs, but also the concentrations of the paralogous TFs, the presence and concentrations of co-factor proteins^107, 115–117^, the chromatin environment, etc. Our study takes an important first step in deciphering the molecular mechanisms of differential specificity in TF families, by identifying differences in intrinsic preferences between paralogous TFs and showing that these *in vitro* differences partially explain differential *in vivo* binding. We envision that more quantitative high-throughput technologies and computational models will be developed to gain a deeper understanding of the differential genomic binding and function of paralogous TFs.

## METHODS [online]

Methods and associated references are available as Supplementary Material.

## 5 AUTHOR CONTRIBUTIONS

NS, RG designed the research. NS, JS, TB, JH designed and performed experiments. NS, YZ performed data analysis. NS, JZ carried out computational modeling. DL, JB, HL implemented the web server. NS, JZ, and RG wrote the manuscript with the participation of all authors.

## ACKNOWLEGMENTS

This work was funded by NSF grant MCB-14-12045, NIH grant R01-GM117106, and an Alfred P. Sloan Foundation fellowship (to RG). High-performance computing was partially supported through North Carolina Biotechnology Center grant 2016-IDG-1013 (to HL). The melanoma data used in this study was generated as part of the SKCA-BR project, funded by Barretos Cancer Hospital (Brazil). The authors thank members of the Gordan lab, Dr. Jen-Tsan Ashley Chi, Dr. Andrew Allen, Dr. Brendan Frey, and members of the Frey group for helpful discussions and feedback.

## REFERENCES

1. Ayer, D.E., Kretzner, L. & Eisenman, R.N. Mad: a heterodimeric partner for Max that antagonizes Myc transcriptional activity. Cell 72, 211–222 (1993).

2. Yadav, S., Mukhopadhyay, S., Anbalagan, M. & Makridakis, N. Somatic Mutations in Catalytic Core of POLK Reported in Prostate Cancer Alter Translesion DNA Synthesis. Hum Mutat 36, 873–880(2015).

3. International Cancer Genome, C. et al. International network of cancer genome projects. Nature 464, 993–998 (2010).

4. Rose, P.W. et al. The RCSB Protein Data Bank: views of structural biology for basic and applied research and education. Nucleic Acids Res 43, D345–356 (2015).

5. Best, D.J. & Roberts, D.E. Algorithm AS 89: The Upper Tail Probabilities of Spearman's rho. Applied Statistics 24, 377–379 (1975.

6. Genomes Project, C. et al. An integrated map of genetic variation from 1,092 human genomes. Nature 491, 56–65 (2012.

7. Zhou, J. & Troyanskaya, O.G. Predicting effects of noncoding variants with deep learning-based sequence model. Nat Methods 12, 931–934 (2015).

8. Zhang, J. Evolution by gene duplication: an update. Trends in ecology & evolution 18, 292–298 (2003).

9. Cheatle Jarvela, A.M. & Hinman, V.F. Evolution of transcription factor function as a mechanism for changing metazoan developmental gene regulatory networks. Evodevo 6, 3 (2015).

10. Chen, K. & Rajewsky, N. The evolution of gene regulation by transcription factors and microRNAs. Nat Rev Genet 8, 93–103 (2007).

11. Hsia, C.C. & McGinnis, W. Evolution of transcription factor function. Curr Opin Genet Dev 13, 199–206 (2003).

12. Hughes, A.L. The evolution of functionally novel proteins after gene duplication. Proc Biol Sci 256, 119–124 (1994).

13. Lynch, M. & Conery, J.S. The evolutionary fate and consequences of duplicate genes. Science 290, 1151–1155 (2000).

14. Taylor, J.S. & Raes, J. Duplication and divergence: the evolution of new genes and old ideas. Annu Rev Genet 38, 615–643 (2004).

15. Wagner, A. Selection and gene duplication: a view from the genome. Genome Biol 3, reviews1012 (2002).

16. Vaquerizas, J.M., Kummerfeld, S.K., Teichmann, S.A. & Luscombe, N.M. A census of human transcription factors: function, expression and evolution. Nat Rev Genet 10, 252–263 (2009).

17. Landt, S.G. et al. ChIP-seq guidelines and practices of the ENCODE and modENCODE consortia. Genome Res 22, 1813–1831 (2012).

18. Graves, B.J. & Petersen, J.M. Specificity within the ets family of transcription factors. Adv Cancer Res 75, 1–55 (1998).

19. Hollenhorst, P.C., McIntosh, L.P. & Graves, B.J. Genomic and biochemical insights into the specificity of ETS transcription factors. Annu Rev Biochem 80, 437–471 (2011).

20. Wei, G.H. et al. Genome-wide analysis of ETS-family DNA-binding in vitro and in vivo. The EMBO journal 29, 2147–2160 (2010).

21. Cesari, F. et al. Mice deficient for the ets transcription factor elk-1 show normal immune responses and mildly impaired neuronal gene activation. Mol Cell Biol 24, 294–305 (2004).

22. Eyquem, S., Chemin, K., Fasseu, M. & Bories, J.C. The Ets-1 transcription factor is required for complete pre-T cell receptor function and allelic exclusion at the T cell receptor beta locus. Proceedings of the National Academy of Sciences of the United States of America 101, 15712–15717 (2004).

23. Eyquem, S. et al. The development of early and mature B cells is impaired in mice deficient for the Ets-1 transcription factor. Eur J Immunol 34, 3187–3196 (2004).

24. Barton, K. et al. The Ets-1 transcription factor is required for the development of natural killer cells in mice. Immunity 9, 555–563 (1998).

25. Gallant, S. & Gilkeson, G. ETS transcription factors and regulation of immunity. Arch Immunol Ther Exp (Warsz) 54, 149–163 (2006).

26. Mouly, E. et al. The Ets-1 transcription factor controls the development and function of natural regulatory T cells. J Exp Med 207, 2113–2125 (2010).

27. Besnard, A., Galan-Rodriguez, B., Vanhoutte, P. & Caboche, J. Elk-1 a transcription factor with multiple facets in the brain. Front Neurosci 5, 35 (2011).

28. Boros, J. et al. Overlapping promoter targeting by Elk-1 and other divergent ETS-domain transcription factor family members. Nucleic Acids Res 37, 7368–7380 (2009).

29. Grinkevich, L.N., Lisachev, P.D., Gudzik, K.A., Grinkevich, V.V. & Kharchenko, O.A. Comparative analysis of the activation of the Elk-1 transcription factor in the central nervous system of animals with different learning capacities. Dokl Biol Sci 397, 269–271 (2004).

30. Boros, J. et al. Elucidation of the ELK1 target gene network reveals a role in the coordinate regulation of core components of the gene regulation machinery. Genome research 19, 1963–1973 (2009).

31. Li, Q.J., Vaingankar, S., Sladek, F.M. & Martins-Green, M. Novel nuclear target for thrombin: activation of the Elk1 transcription factor leads to chemokine gene expression. Blood 96, 3696–3706 (2000).

32. Odrowaz, Z. & Sharrocks, A.D. ELK1 uses different DNA binding modes to regulate functionally distinct classes of target genes. PLoS Genet 8, e1002694 (2012.

33. Patki, M. et al. The ETS domain transcription factor ELK1 directs a critical component of growth signaling by the androgen receptor in prostate cancer cells. The Journal of biological chemistry 288, 11047-11065 (2013).

34. Weirauch, M.T. et al. Determination and inference of eukaryotic transcription factor sequence specificity. Cell 158, 1431–1443 (2014).

35. Badis, G. et al. Diversity and complexity in DNA recognition by transcription factors. Science 324, 1720-1723 (2009).

36. Weirauch, M.T. et al. Evaluation of methods for modeling transcription factor sequence specificity. Nature biotechnology 31, 126–134 (2013).

37. Jolma, A. et al. DNA-binding specificities of human transcription factors. Cell 152, 327–339 (2013).

38. Jolma, A. et al. Multiplexed massively parallel SELEX for characterization of human transcription factor binding specificities. Genome research 20, 861–873 (2010).

39. Alipanahi, B., Delong, A., Weirauch, M.T. & Frey, B.J. Predicting the sequence specificities of DNA- and RNA-binding proteins by deep learning. Nature biotechnology 33, 831–838 (2015).

40. Wang, J. et al. Factorbook.org: a Wiki-based database for transcription factor-binding data generated by the ENCODE consortium. Nucleic Acids Res 41, D171–176 (2013).

41. Foley, K.P. & Eisenman, R.N. Two MAD tails: what the recent knockouts of Mad1 and Mxi1 tell us about the MYC/MAX/MAD network. Biochim Biophys Acta 1423, M37–47 (1999).

42. Meyer, N. & Penn, L.Z. Reflecting on 25 years with MYC. Nat Rev Cancer 8, 976–990 (2008).

43. Dang, C.V. MYC on the path to cancer. Cell 149, 22–35 (2012).

44. Johnson, D.S., Mortazavi, A., Myers, R.M. & Wold, B. Genome-wide mapping of in vivo protein-DNA interactions. Science 316, 1497–1502 (2007).

45. Gordan, R. et al. Genomic regions flanking E-box binding sites influence DNA binding specificity of bHLH transcription factors through DNA shape. Cell Rep 3, 1093–1104 (2013).

46. Consortium, E.P. An integrated encyclopedia of DNA elements in the human genome. Nature 489, 57–74 (2012).

47. Zervos, A.S., Gyuris, J. & Brent, R. Mxi1, a protein that specifically interacts with Max to bind Myc-Max recognition sites. Cell 72, 223–232 (1993).

48. Matys, V. et al. TRANSFAC and its module TRANSCompel: transcriptional gene regulation in eukaryotes. Nucleic Acids Res 34, D108–110 (2006).

49. Berger, M.F. et al. Compact, universal DNA microarrays to comprehensively determine transcription-factor binding site specificities. Nat Biotechnol 24, 1429–1435 (2006).

50. Berger, M.F. & Bulyk, M.L. Universal protein-binding microarrays for the comprehensive characterization of the DNA-binding specificities of transcription factors. Nature protocols 4, 393–411 (2009).

51. Annala, M., Laurila, K., Lahdesmaki, H. & Nykter, M. A linear model for transcription factor binding affinity prediction in protein binding microarrays. PloS one 6, e20059 (2011).

52. Komori, T. Regulation of bone development and maintenance by Runx2. Front Biosci 13, 898–903 (2008).

53. Elagib, K.E. et al. RUNX1 and GATA-1 coexpression and cooperation in megakaryocytic differentiation. Blood 101, 4333–4341 (2003).

54. Lacaud, G. et al. Runx1 is essential for hematopoietic commitment at the hemangioblast stage of development in vitro. Blood 100, 458–466 (2002).

55. Hyde, R.K., Zhao, L., Alemu, L. & Liu, P.P. Runx1 is required for hematopoietic defects and leukemogenesis in Cbfb-MYH11 knock-in mice. Leukemia 29, 1771–1778 (2015).

56. Komori, T. Signaling networks in RUNX2-dependent bone development. J Cell Biochem 112, 750–755 (2011).

57. Liu, T.M. & Lee, E.H. Transcriptional regulatory cascades in Runx2-dependent bone development. Tissue Eng Part B Rev 19, 254–263 (2013).

58. Siggers, T. & Gordan, R. Protein-DNA binding: complexities and multi-protein codes. Nucleic Acids Res 42, 2099–2111 (2014).

59. Scardigli, R., Baumer, N., Gruss, P., Guillemot, F. & Le Roux, I. Direct and concentration-dependent regulation of the proneural gene Neurogenin2 by Pax6. Development 130, 3269–3281 (2003).

60. Gaudet, J. & Mango, S.E. Regulation of organogenesis by the Caenorhabditis elegans FoxA protein PHA-4. Science 295, 821–825 (2002).

61. Jaeger, S.A. et al. Conservation and regulatory associations of a wide affinity range of mouse transcription factor binding sites. Genomics 95, 185–195 (2010).

62. Tanay, A. Extensive low-affinity transcriptional interactions in the yeast genome. Genome research 16, 962–972 (2006).

63. Drucker, H., Burges, C.J.C., Kaufman, L., Smola, A. & Vapnik, V. Support vector regression machines. Advances in Neural Information Processing Systems 9, 155–161 (1997).

64. Zhou, T. et al. Quantitative modeling of transcription factor binding specificities using DNA shape. Proceedings of the National Academy of Sciences of the United States of America 112, 4654–4659 (2015).

65. Mordelet, F., Horton, J., Hartemink, A.J., Engelhardt, B.E. & Gordan, R. Stability selection for regression-based models of transcription factor-DNA binding specificity. Bioinformatics 29,i117–125 (2013).

66. Yang, L. et al. TFBSshape: a motif database for DNA shape features of transcription factor binding sites. Nucleic Acids Res 42, D148–155 (2014).

67. Rohs, R. et al. Origins of specificity in protein-DNA recognition. Annu Rev Biochem 79, 233–269 (2010).

68. Zhou, T. et al. DNAshape: a method for the high-throughput prediction of DNA structural features on a genomic scale. Nucleic Acids Res 41, W56–62 (2013).

69. Kidder, B.L., Hu, G. & Zhao, K. ChIP-Seq: technical considerations for obtaining high-quality data. Nat Immunol 12, 918–922 (2011).

70. Roeder, R.G. The role of general initiation factors in transcription by RNA polymerase II. Trends Biochem Sci 21, 327–335 (1996).

71. Poorey, K. et al. Measuring chromatin interaction dynamics on the second time scale at singlecopy genes. Science 342, 369–372 (2013).

72. Solomon, M.J. & Varshavsky, A. Formaldehyde-mediated DNA-protein crosslinking: a probe for in vivo chromatin structures. Proceedings of the National Academy of Sciences of the United States of America 82, 6470–6474 (198).

73. Hussain Zaidi, E.A.H., Savera J. Shetty, Stefan Bekiranov, David T. Auble Second-generation method for analysis of chromatin binding using formaldehyde crosslinking kinetics. bioRxiv.

74. Lu, K. et al. Structural characterization of formaldehyde-induced cross-links between amino acids and deoxynucleosides and their oligomers. J Am Chem Soc 132, 3388–3399 (2010).

75. Gavrilov, A., Razin, S.V. & Cavalli, G. In vivo formaldehyde cross-linking: it is time for black box analysis. Briefings in functional genomics 14, 163–165 (2015).

76. Parseghian, M.H. Hitchhiker antigens: inconsistent ChIP results, questionable immunohistology data, and poor antibody performance may have a common factor. Biochem Cell Biol 91, 378–394 (2013).

77. Schonbrunn, A. Editorial: Antibody can get it right: confronting problems of antibody specificity and irreproducibility. Mol Endocrinol 28, 1403–1407 (2014).

78. Wardle, F.C. & Tan, H. A ChIP on the shoulder? Chromatin immunoprecipitation and validation strategies for ChIP antibodies. F1000Res 4, 235 (2015).

79. Teytelman, L., Thurtle, D.M., Rine, J. & van Oudenaarden, A. Highly expressed loci are vulnerable to misleading ChIP localization of multiple unrelated proteins. Proceedings of the National Academy of Sciences of the United States of America 110, 18602–18607 (2013).

80. Park, D., Lee, Y., Bhupindersingh, G. & Iyer, V.R. Widespread misinterpretable ChIP-seq bias in yeast. PloS one 8, e83506 (2013).

81. Jain, D., Baldi, S., Zabel, A., Straub, T. & Becker, P.B. Active promoters give rise to false positive ‘Phantom Peaks’ in ChIP-seq experiments. Nucleic Acids Res 43, 6959–6968 (2015).

82. Bardet, A.F., He, Q., Zeitlinger, J. Stark, A. A computational pipeline for comparative ChIP-seq analyses. Nature protocols 7, 45–61 (2011).

83. Poptsova, M.S. et al. Non-random DNA fragmentation in next-generation sequencing. Sci Rep 4, 4532 (2014).

84. Kulakovskiy, I.V. et al. HOCOMOCO: a comprehensive collection of human transcription factor binding sites models. Nucleic Acids Res 41, D195–202 (2013).

85. Kulakovskiy, I.V. et al. HOCOMOCO: expansion and enhancement of the collection of transcription factor binding sites models. Nucleic Acids Res 44, D116–125 (2016).

86. Orenstein, Y. & Shamir, R. A comparative analysis of transcription factor binding models learned from PBM, HT-SELEX and ChIP data. Nucleic Acids Res 42, e63 (2014).

87. Mariani, L., Weinand, K., Vedenko, A., Barrera, L.A. & Bulyk, M.L. Identification of Human Lineage-Specific Transcriptional Coregulators Enabled by a Glossary of Binding Modules and Tunable Genomic Backgrounds. Cell Syst 5, 187–201 e187 (2017).

88. Gordan, R., Hartemink, A.J. & Bulyk, M.L. Distinguishing direct versus indirect transcription factor-DNA interactions. Genome research 19, 2090–2100 (2009).

89. Arvey, A., Agius, P., Noble, W.S. & Leslie, C. Sequence and chromatin determinants of cell-type-specific transcription factor binding. Genome research 22, 1723–1734 (2012).

90. Isakova, A. et al. SMiLE-seq identifies binding motifs of single and dimeric transcription factors. Nat Methods 14, 316–322 (2017).

91. Mahony, S. et al. An integrated model of multiple-condition ChIP-Seq data reveals predeterminants of Cdx2 binding. PLoS computational biology 10, e1003501 (2014).

92. Tuteja, G., White, P., Schug, J. & Kaestner, K.H. Extracting transcription factor targets from ChIP-Seq data. Nucleic Acids Res 37, e113 (2009).

93. Kibet, C.K. & Machanick, P. Transcription factor motif quality assessment requires systematic comparative analysis. F1000Res 4 (2015).

94. Gusmao, E.G., Allhoff, M., Zenke, M. & Costa, I.G. Analysis of computational footprinting methods for DNase sequencing experiments. Nat Methods 13, 303–309 (2016).

95. Alberstein, M. et al. Regulation of transcription of the RNA splicing factor hSlu7 by Elk-1 and Sp1 affects alternative splicing. RNA 13, 1988–1999 (2007).

96. Gustafson, W.C. et al. Bcr-Abl regulates protein kinase Ciota (PKCiota) transcription via an Elk1 site in the PKCiota promoter. The Journal of biological chemistry 279, 9400–9408 (2004).

97. Fukuda, M., Gotoh, Y. & Nishida, E. Interaction of MAP kinase with MAP kinase kinase: its possible role in the control of nucleocytoplasmic transport of MAP kinase. The EMBO journal 16, 1901–1908 (1997).

98. Bories, J.C. et al. Increased T-cell apoptosis and terminal B-cell differentiation induced by inactivation of the Ets-1 proto-oncogene. Nature 377, 635–638 (1995).

99. Teruyama, K. et al. Role of transcription factor Ets-1 in the apoptosis of human vascular endothelial cells. J Cell Physiol 188, 243–252 (2001).

100. Hofmann, T.G. & Will, H. Body language: the function of PML nuclear bodies in apoptosis regulation. Cell Death Differ 10, 1290–1299 (2003).

101. Soldatenkov, V.A. et al. Differential regulation of the response to DNA damage in Ewing's sarcoma cells by ETS1 and EWS/FLI-1. Oncogene 21, 2890–2895 (2002).

102. Andersen, M.C. et al. In silico detection of sequence variations modifying transcriptional regulation. PLoS computational biology 4, e5 (2008).

103. Thomas-Chollier, M. et al. Transcription factor binding predictions using TRAP for the analysis of ChIP-seq data and regulatory SNPs. Nature protocols 6, 1860–1869 (2011).

104. Ward, L.D. & Kellis, M. HaploReg v4: systematic mining of putative causal variants, cell types, regulators and target genes for human complex traits and disease. Nucleic Acids Res 44, D877–881 (2016).

105. McVicker, G. et al. Identification of genetic variants that affect histone modifications in human cells. Science 342, 747–749 (2013).

106. Lee, D. et al. A method to predict the impact of regulatory variants from DNA sequence. Nat Genet 47, 955–961 (2015).

107. Slattery, M. et al. Cofactor binding evokes latent differences in DNA binding specificity between Hox proteins. Cell 147, 1270–1282 (2011).

108. Orlando, D.A. et al. Quantitative ChIP-Seq normalization reveals global modulation of the epigenome. Cell Rep 9, 1163–1170 (2014).

109. Grzybowski, A.T., Chen, Z. & Ruthenburg, A.J. Calibrating ChIP-Seq with Nucleosomal Internal Standards to Measure Histone Modification Density Genome Wide. Mol Cell 58, 886–899 (2015).

110. van Galen, P. et al. A Multiplexed System for Quantitative Comparisons of Chromatin Landscapes. Mol Cell 61, 170–180 (2016).

111. Skene, P.J. & Henikoff, S. An efficient targeted nuclease strategy for high-resolution mapping of DNA binding sites. Elife 6 (2017).

112. Zentner, G.E., Kasinathan, S., Xin, B., Rohs, R. & Henikoff, S. ChEC-seq kinetics discriminates transcription factor binding sites by DNA sequence and shape in vivo. Nat Commun 6, 8733 (2015).

113. Rhee, H.S. & Pugh, B.F. Comprehensive genome-wide protein-DNA interactions detected at single-nucleotide resolution. Cell 147, 1408–1419 (2011).

114. He, Q., Johnston, J. & Zeitlinger, J. ChIP-nexus enables improved detection of in vivo transcription factor binding footprints. Nature biotechnology 33, 395–401 (2015).

115. Siggers, T., Duyzend, M.H., Reddy, J., Khan, S. & Bulyk, M.L. Non-DNA-binding cofactors enhance DNA-binding specificity of a transcriptional regulatory complex. Mol Syst Biol 7, 555 (2011).

116. Mann, R.S., Lelli, K.M. & Joshi, R. Hox specificity unique roles for cofactors and collaborators. Curr Top Dev Biol 88, 63–101 (2009).

117. Garvie, C.W., Hagman, J. & Wolberger, C. Structural studies of Ets-1/Pax5 complex formation on DNA. Molecular Cell 8, 1267–1276 (2001).

